## Supplementary information for "Noise-augmented directional clustering of genetic association data identifies distinct mechanisms underlying obesity"

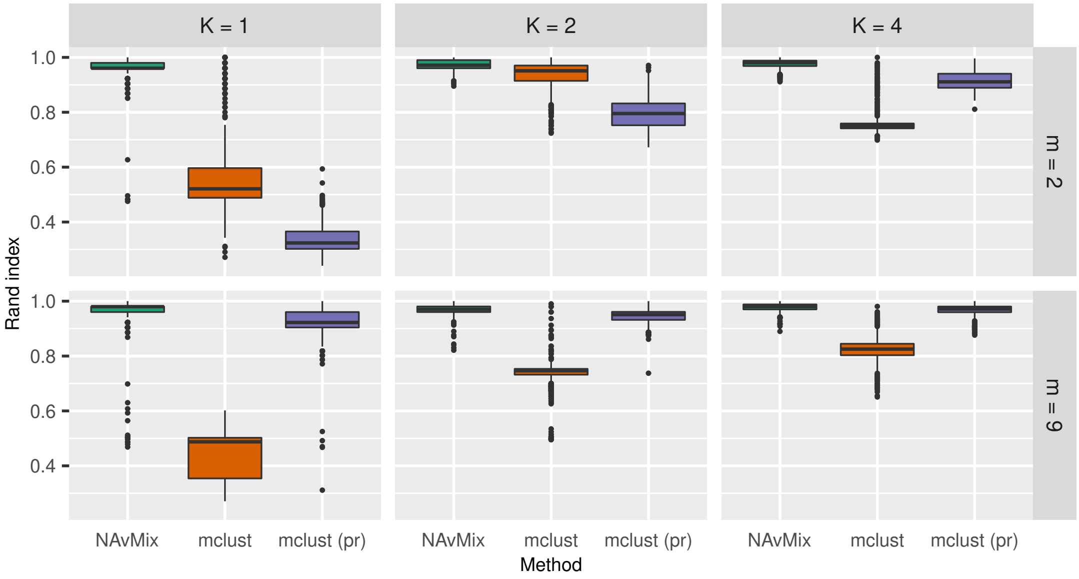


Figure S1: Boxplots of the Rand index for each scenario using NAvMix, mclust, and mclust with proportional effects (pr) when the estimated trait correlation was incorporated.

Table S1: Mean number of clusters estimated and mean number of observations allocated to the noise cluster for each simulated scenario using NAvMix, mclust, and mclust using proportional associations (pr) when the estimated trait correlation was incorporated.

| Number of traits ($m$) | Number of clusters ($K$) | Number of clusters | | | Number of noise variants | | |
|---|---|---|---|---|---|---|---|
| | | NAvMix | mclust | mclust (pr) | NAvMix | mclust | mclust (pr) |
| 2 | 1 | 1.01 | 1.98 | 3.89 | 19.01 | 15.13 | 19.05 |
| | 2 | 2.00 | 2.09 | 4.63 | 17.33 | 12.24 | 17.71 |
| | 4 | 4.00 | 2.15 | 6.16 | 14.21 | 14.21 | 14.99 |
| 9 | 1 | 1.02 | 2.50 | 1.03 | 21.45 | 18.32 | 23.70 |
| | 2 | 2.01 | 4.65 | 2.01 | 23.26 | 16.03 | 25.45 |
| | 4 | 4.01 | 5.97 | 4.52 | 23.33 | 18.08 | 23.61 |

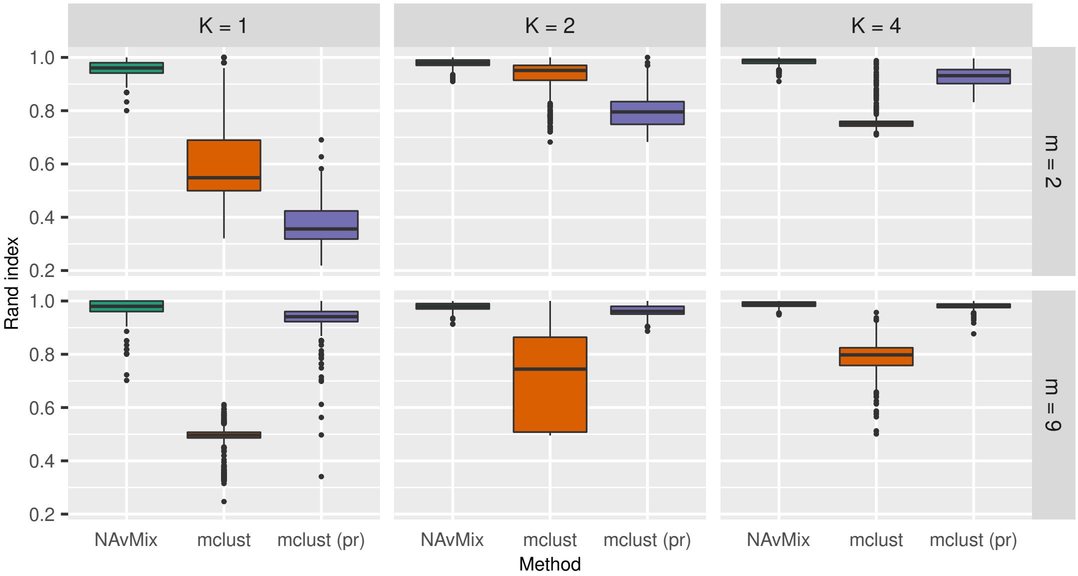


Figure S2: Boxplots of the Rand index for each scenario using NAvMix, mclust, and mclust with proportional effects (pr) when the genetic variant-trait associations were estimated in difference samples of different size.

Table S2: Mean number of clusters estimated and mean number of observations allocated to the noise cluster for each simulated scenario using NAvMix, mclust, and mclust using proportional associations (pr) when the genetic variant-trait associations were estimated in difference samples of different size.

| Number of traits ($m$) | Number of clusters ($K$) | Number of clusters | | | Number of noise variants | | |
|---|---|---|---|---|---|---|---|
| | | NAvMix | mclust | mclust (pr) | NAvMix | mclust | mclust (pr) |
| 2 | 1.00 | 1.00 | 1.86 | 3.72 | 18.91 | 14.18 | 19.15 |
| | 2 | 2.00 | 2.16 | 4.67 | 17.06 | 12.63 | 16.97 |
| | 4 | 4.00 | 2.13 | 6.14 | 12.36 | 14.83 | 14.02 |
| 9 | 1 | 1.01 | 2.12 | 1.05 | 21.27 | 17.86 | 22.92 |
| | 2 | 2.00 | 2.98 | 2.00 | 22.19 | 17.73 | 23.73 |
| | 4 | 4.87 | 4.39 | 4.98 | 17.78 | 18.51 | 22.04 |

Table S4: Intercept estimate, standard error of estimate (SE), 95% confidence interval (CI) and p-value from the MR-Egger intercept test when examining the association of genetically predicted BMI with CHD, for all genetic variants and for each cluster.

| Instruments | Estimate | SE | CI | p-value |
|---|---|---|---|---|
| All genetic variants | -0.002 | 0.002 | (-0.006, 0.001) | 0.135 |
| Cluster 1 | -0.002 | 0.002 | (-0.006, 0.002) | 0.335 |
| Cluster 2 | 0.002 | 0.002 | (-0.002, 0.007) | 0.358 |
| Cluster 3 | -0.002 | 0.005 | (-0.011, 0.007) | 0.657 |
| Cluster 4 | 0.006 | 0.007 | (-0.009, 0.020) | 0.442 |
| Cluster 5 | -0.022 | 0.015 | (-0.050, 0.007) | 0.133 |

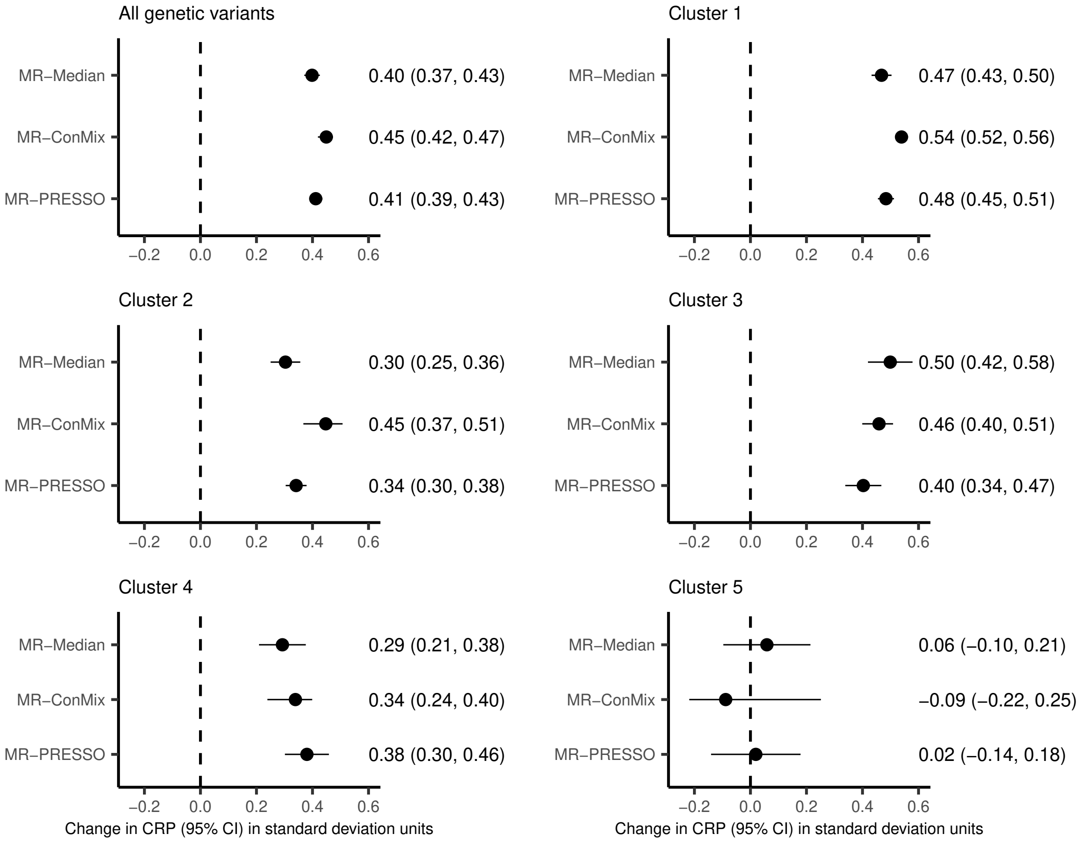


Figure S3: Estimates and 95% confidence intervals of the association of genetically predicted BMI on CRP from MR-Median, the Contamination Mixture method (MR-ConMix) and MR-PRESSO, for all genetic variants and for each cluster. Estimates represent the change in CRP in standard deviation units per 1 standard deviation increase in genetically predicted BMI.
